# Markers of Fungal Translocation Are Elevated During Post-Acute Sequelae of SARS-CoV-2 Infection and Induce NF-κB Triggered Inflammation

**DOI:** 10.1101/2022.04.12.488051

**Authors:** Leila B. Giron, Michael J. Peluso, Jianyi Ding, Grace Kenny, Netanel F Zilberstein, Jane Koshy, Kai Ying Hong, Heather Rasmussen, Greg Miller, Faraz Bishehsari, Robert A. Balk, James N. Moy, Rebecca Hoh, Scott Lu, Aaron R. Goldman, Hsin-Yao Tang, Brandon C. Yee, Ahmed Chenna, John W. Winslow, Christos J. Petropoulos, J. Daniel Kelly, Haimanot Wasse, Jeffrey N. Martin, Qin Liu, Ali Keshavarzian, Alan Landay, Steven G. Deeks, Timothy J. Henrich, Mohamed Abdel-Mohsen

## Abstract

Long COVID, a type of Post-Acute Sequelae of SARS CoV-2 infection (PASC), has been associated with sustained elevated levels of immune activation and inflammation. However, the pathophysiological mechanisms that drive this inflammation remain unknown. Inflammation during acute Coronavirus Disease 2019 (COVID-19) could be exacerbated by microbial translocation (from the gut and/or lung) to the blood. Whether microbial translocation contributes to inflammation during PASC is unknown. We found higher levels of fungal translocation – measured as β-glucan, a fungal cell wall polysaccharide – in the plasma of individuals experiencing PASC compared to those without PASC or SARS-CoV-2 negative controls. The higher β-glucan correlated with higher levels of markers of inflammation and elevated levels of host metabolites involved in activating *N*-Methyl-D-aspartate receptors (such as metabolites within the tryptophan catabolism pathway) with established neuro-toxic properties. Mechanistically, β-glucan can directly induce inflammation by binding to myeloid cells (via the Dectin-1 receptor) and activating Syk/NF-κB signaling. Using an *in vitro* Dectin-1/NF-κB reporter model, we found that plasma from individuals experiencing PASC induced higher NF-κB signaling compared to plasma from SARS-CoV-2 negative controls. This higher NF-κB signaling was abrogated by the Syk inhibitor Piceatannol. These data suggest a potential targetable mechanism linking fungal translocation and inflammation during PASC.

## INTRODUCTION

SARS-CoV-2 infection causes acute respiratory and systemic disease (COVID-19) (1, 2). A subset of individuals also experience persistent, recurrent, or new COVID-19-attributed symptoms in the months following acute infection – a condition commonly referred to as Long COVID (3–19). Long COVID, a type of Post-Acute Sequelae of SARS CoV-2 infection (PASC), can impact an individual’s overall health and quality of life (3–18, 20). Recently, PASC has been associated with sustained elevated levels of immune activation and inflammation (21–24). However, the pathophysiological mechanisms that drive this inflammation remain unknown. Among the hypothesized drivers are pre-existing medical comorbidities, such as diabetes or obesity, the degree of SARS-CoV-2 viremia during acute infection, latent Epstein-Barr virus reactivation, and the production of autoantibodies (25–27). We have been investigating a known driver of systemic inflammation and severity during other respiratory-related diseases, microbial translocation resulting from disruption in the gut-lung axis.

Disruption of the gut-lung axis is a known marker of severity during other respiratory diseases (28–31) and may play a role in potentiating worse clinical outcomes. SARS-CoV-2 infection can affect the gastrointestinal tract (GI) tract and cause GI symptoms (32, 33) through indirect and/or direct mechanisms. Indirectly, lung infection or injury can provoke systemic inflammation (including cytokine storm), which in turn can disrupt gut barrier integrity (mainly by IFNγ and TNFα which are known to disrupt tight junction permeability (34–36)), enabling gut microbes and their products to translocate across the gut epithelium. This translocation (which can also happen across the lung epithelium) can exacerbate the initial systemic inflammation, resulting in a positive feedback loop (28-31, 37-39). Directly, SARS-CoV-2 can infect gut cells (40); other viral infections of the gut can cause a breakdown of the epithelial barrier (41–43).

Acute COVID-19 has been associated with an increase in the plasma levels of zonulin, an established physiological driver of tight junction permeability (44, 45). This increased permeability enables the translocation of both bacterial and fungal products to the blood. Such microbial translocation correlates with increased systemic inflammation, disrupted gut-associated metabolites, and higher mortality during acute COVID-19 (46). These observations are supported with a series of recent studies, using stool samples, showing that COVID-19 severity is associated with a state of gut microbial dysbiosis and translocation (including fungal translocation) (47–54). Although these data (46–53) do not imply that gut microbial translocation is the primary trigger of inflammation during COVID-19, as the clinical syndrome of COVID-19 likely embodies multiple pathophysiological pathways, they are consistent with the literature indicating that microbial translocation fuels inflammation and disease severity during respiratory diseases (28–31), and thus support a model in which microbial translocation fuels inflammation following SARS-CoV-2 infection. However, whether the translocation of microbes –bacteria or fungus – is related to inflammation during PASC is unknown and is the subject of this study.

## RESULTS

### Participant characteristics

We used samples from two cohorts (**Table 1**): 1) Cross-sectional plasma samples from 117 volunteers with a history of COVID-19 (a subset of the UCSF LIINC cohort) 90-160 days after their first positive SARS-CoV-2 result. These participants were divided into two groups: 56 individuals with no ongoing COVID-19-attributed symptoms at the time of sample collection (non-PASC) and 61 with ≥2 symptoms present at the time of sample collection (PASC; **Table 1**). 2) Cross-sectional plasma samples from 50 COVID-19 individuals experiencing PASC 3-4 months after their convalescence from acute COVID-19 (a subset of the Rush PASC cohort) and cross-sectional plasma samples from 50 SARS-CoV-2 negative controls (matched for age and gender to the Rush PASC samples; **Table 1**).

**Table 1.** Demographic and clinical characteristics of study cohorts

|  |  | UCSF LIINC cohort |  | Rush PASC cohort |  |
| --- | --- | --- | --- | --- | --- |
|  |  | No PASC | PASC | SARS-CoV2 negative | PASC |
| N |  | 56 | 61 | 50 | 50 |
| Age (years; median (IQR)) |  | 49 (19.5) | 49 (19) | 43 (11.8) | 45 (18) |
| Sex (n (%)) | Male | 33 (55%) | 27 (45%) | 19 (38%) | 19 (38) |
|  | Female | 23 (40%) | 34 (60%) | 31 (62%) | 31 (62) |
| Race/ethnicity (n (%)) | Asian | 8 (73%) | 3 (27%) | 0 (0%) | 0 (0%) |
|  | Black | 3 (50%) | 3 (50%) | 26 (52%) | 7 (14) |
|  | Hispanic/Latino | 9 (31%) | 20 (69%) | 5 (10%) | 9 (18) |
|  | White | 31 (48%) | 34 (52%) | 0 (0%) | 34 (68) |
|  | Pacific Islander/Native Hawaiian | 2 (67%) | 1 (33%) | 19 (38%) | 0 (0%) |
| Pre-existing Comorbidities (autoimmune disease, cancer, diabetes, hypertension, heart disease, lung disease, kidney disease, liver disease) (n (%)) | No | 31 (51%) | 30 (49%) | - | - |
|  | Yes | 25 (45%) | 31 (55%) | - | - |
| BMI (Median (IQR)) |  | 26.14 (7.25) | 30.94 (9.9) | 34.9 (4.9) | 28.7 (7.2) |
| Number of symptoms (Median (IQR)) |  | 0 (0) | 6 (4) | 0 (0) | 4 (3) |
| Hospitalization during acute COVID-19 illness (n (%)) | Nonhospitalized | 47 (51%) | 46 (49%) | - | 38 (76) |
|  | Hospitalized | 9 (37.5%) | 15 (62.5%) | - | 12 (24) |

We examined whether age (**Fig. 1A**), body mass index (BMI; **Fig. 1B**), self-rated overall health/quality of life (QoL) score on a visual-analog scale (0-100; **Fig. 1C**), gender, ethnicity, hospitalization during acute COVID-19, or pre-existing co-morbidities (**Fig. 1D**) differentiate PASC from non-PASC groups within the 117 samples from the UCSF LIINC cohort. We found that a higher BMI (*P*= 0.006; **Fig. 1B**) and a higher rate of pre-existing hypertension (*P*= 0.003; **Fig. 1D**) were associated with the PASC phenotype. The overall health/QoL score was lower in volunteers experiencing PASC than those in the non-PASC group (*P*< 0.0001; **Fig. 1C**). Based on these observations, we adjusted our subsequent analyses on the potential role of microbial translocation in PASC for BMI and hypertension as potential confounders of the PASC phenotype in this subset from the UCSF LIINC cohort. We also used the overall health/QoL score in our subsequent analyses examining the potential impact of microbial translocation on individuals’ well-being during PASC.

**Figure 1.**
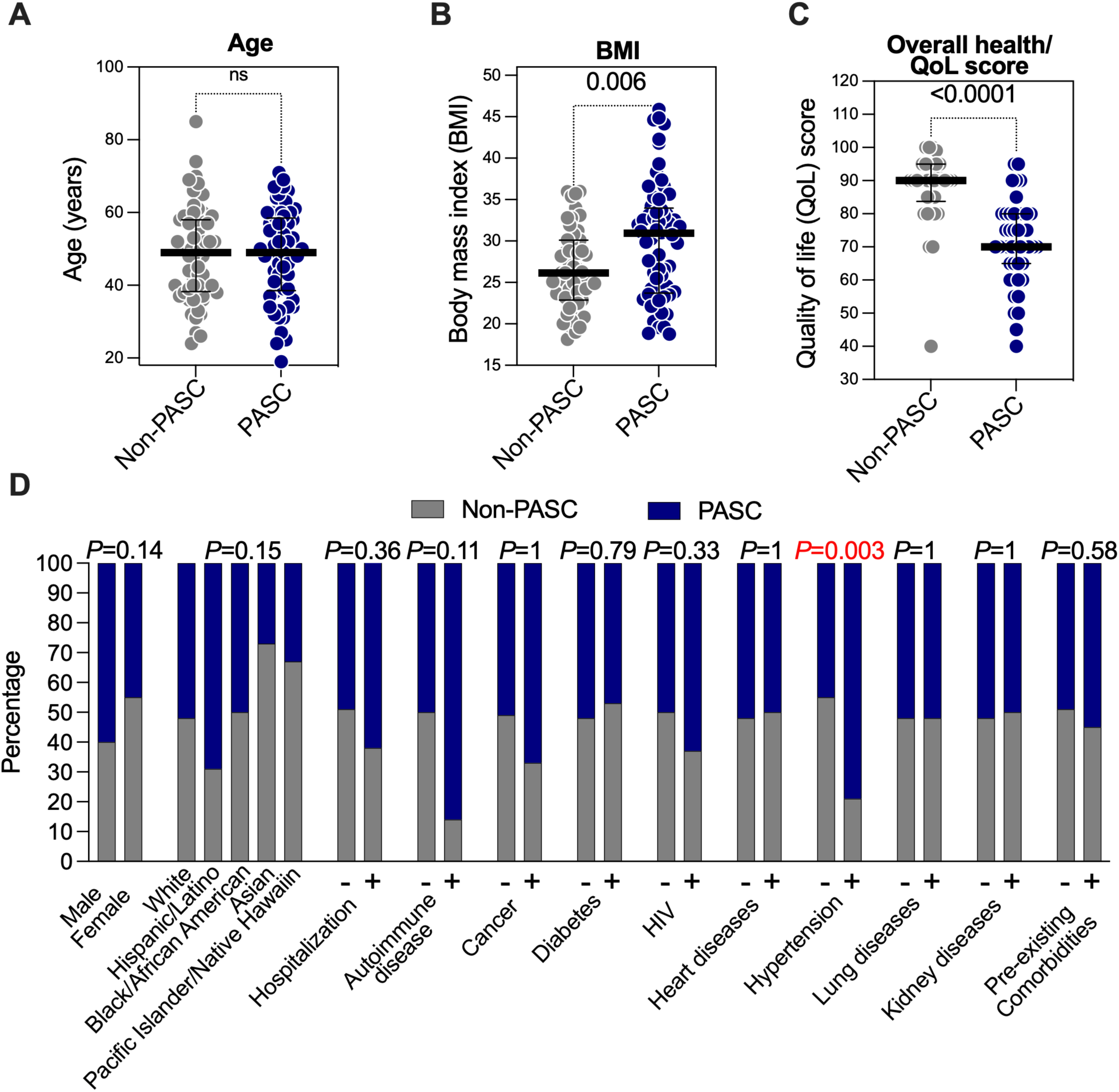
Body mass index (BMI) and hypertension differentiate PASC from non-PASC in a subset of the UCSF LIINC cohort. Demographic and clinical characteristics of 117 individuals from the UCSF LIINC cohort. Out of these 117 individuals, 61 individuals were experiencing two or more COVID-19-attributed symptoms four months following SARS-CoV-2 infection (PASC), whereas 56 individuals were not experiencing any ongoing symptoms (non-PASC). **(A-C)** Mann– Whitney U comparisons of **(A)** age, **(B)** BMI, and **(C)** overall health/quality of life (QoL) score between PASC and non-PASC within the 117 individuals from the UCSF LIINC cohort. Median and interquartile range (IQR) are displayed. **(D)** Fisher’s exact comparisons of the indicated demographic and clinical characteristics between PASC and non-PASC within the 117 individuals from the UCSF LIINC cohort. + = yes and - = no.

### PASC is associated with elevated levels of fungal translocation independent of BMI and hypertension

We first examined levels of tight junction permeability (measured as plasma levels of zonulin) in the plasma of the 117 volunteers from the UCSF LIINC cohort. Zonulin is an established physiological mediator of tight junction permeability in the digestive tract, where higher levels of zonulin drive an increase in fungal and bacterial translocation (44, 55, 56). We found that PASC is associated with an increase in the plasma levels of zonulin compared to non-PASC **(Fig. 2A)**. We next examined levels of fungal translocation (measured as β-glucan, a fungal wall polysaccharide). We observed higher levels of β-glucan in the plasma of volunteers with PASC than non-PASC volunteers (in a manner linked to the number of persistent symptoms and regardless of whether volunteers had been outpatients or hospitalized during their acute COVID-19; **Fig. 2B-E**). Recently, it was shown that plasma β-glucan levels ≥40 pg/ml are clinically significant and associated with higher inflammation and worse survival in patients with acute respiratory failure (57). In the UCSF LIINC cohort, 33% of volunteers experiencing PASC had β-glucan levels ≥40 pg/ml, whereas only 7.1% of non-PASC volunteers had β-glucan ≥40 pg/ml. We also divided the PASC to three PASC phenotypes based on clinical symptom clusters, defined as having at least one symptom in the cluster **–** gastrointestinal (nausea, diarrhea, loss of appetite, abdominal pain, vomiting), cardiopulmonary (cough, dyspnea, chest pain, palpitations), and neurocognitive (headache, concentration problems, dizziness, balance problems, neuropathy, vision problems). β-glucan levels were higher in individuals experiencing each of the three PASC symptom clusters compared to individuals who are not experiencing PASC (**Fig. 2F-H**). Furthermore, we investigated individuals experiencing each symptom separately and found that β-glucan levels were higher in individuals suffering from certain symptoms such as gastrointestinal (GI) symptoms (nausea and diarrhea) as well as vision problems, sleep problems, neuropathy, and pain **(Supplementary Fig. 1)**. We finally examined levels of bacterial translocation (measured as lipopolysaccharide binding protein (LBP)); these levels were not significantly different between the two groups, albeit a trend of higher levels of LBP in individuals experiencing PASC (than non-PASC) was observed **(Fig. 2I)**. In addition, levels of soluble sCD14 and soluble sCD163 (markers of microbial-mediated myeloid inflammation) were not significantly different between the two groups (*P*>0.05).

**Figure 2.**
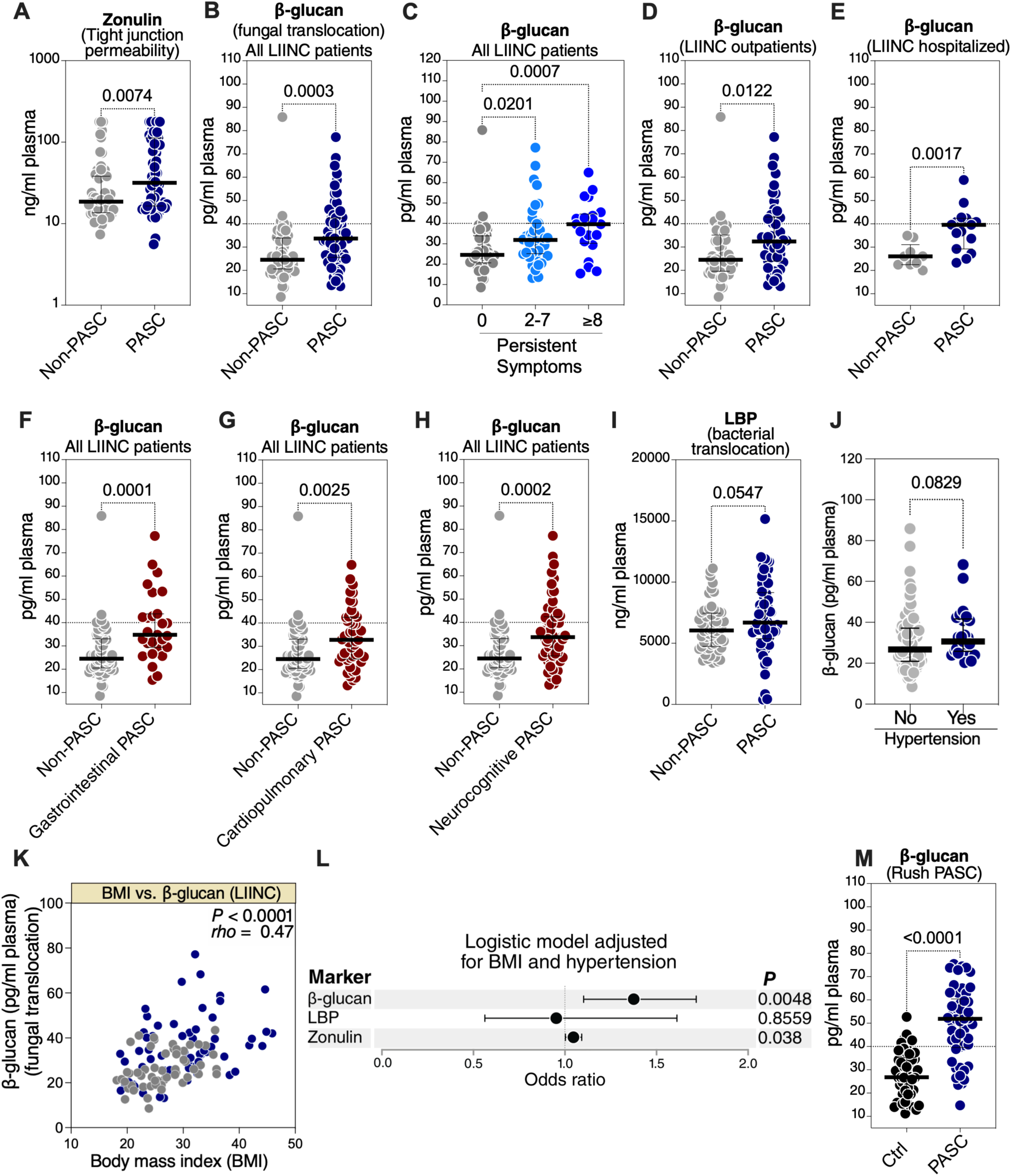
PASC is associated with elevated levels of plasma makers of fungal translocation. **(A)** Levels of zonulin in the plasma of 117 individuals from the UCSF LIINC cohort. Median and IQR are displayed. **(B-E)** Plasma levels of β-glucan in the plasma of 117 individuals from the UCSF LIINC cohort. β-glucan levels are higher in PASC compared to non-PASC when analyzing all individuals (B), dividing the PASC group into individuals with 2-7 symptoms (n=40) or ≥8 symptoms (n=21) (C), analyzing only samples from individuals that were managed as outpatients during their acute COVID-19 illness (D), or analyzing only samples from individuals hospitalized during their acute COVID-19 illness (E). Mann–Whitney U tests. Median and IQR are displayed. **(F-H)** PASC was divided to three PASC phenotypes based on clinical symptom clusters, defined as having at least one symptom in the cluster **–** gastrointestinal (nausea, diarrhea, loss of appetite, abdominal pain, vomiting), cardiopulmonary (cough, dyspnea, chest pain, palpitations), and neurocognitive (headache, concentration problems, dizziness, balance problems, neuropathy, vision problems). β-glucan levels were higher in individuals experiencing each of the three PASC symptom clusters compared to individuals who are not experiencing PASC. Mann–Whitney U tests. Median and IQR are displayed. **(I)** Levels of LBP in the plasma of 117 individuals from the UCSF LIINC cohort. Median and IQR are displayed. **(J)** Mann–Whitney U comparison of the levels of β-glucan in individuals with or without a history of hypertension within the UCSF LIINC cohort. Median and IQR are displayed. **(K)** Spearman’s rank correlation between BMI and the plasma levels of β-glucan in the UCSF LIINC cohort. **(L)** A multivariate logistic model showing that the higher levels of β-glucan and zonulin (odds ratio per five units increase) can differentiate PASC from non-PASC within the UCSF LIINC cohort after adjusting for both BMI and hypertension. **(M)** β-glucan in individuals experiencing PASC in an independent validation cohort (Rush PASC cohort) compared to SARS-CoV-2 negative controls (matched for age and gender); Mann–Whitney U test. Median and IQR are displayed.

Given that we identified BMI and hypertension as potential confounders of the PASC phenotype in the UCSF LIINC cohort, we examined whether levels of plasma β-glucan correlate with BMI and/or hypertension. We found that individuals with hypertension tend to have higher levels of β-glucan (**Fig. 2J**). We also found that higher BMI correlates with higher levels of plasma β-glucan (**Fig. 2K**). This is consistent with recent reports suggesting that obesity is associated with changes in the intestinal mycobiome and with increases in levels of plasma β-glucan (58, 59). As such, we used a multivariate logistic regression model adjusting for BMI and hypertension and found that higher levels of β-glucan (odds ratio (OR) 1.4 per every five units increase; *P* = 0.0048) and zonulin (OR 1.05 per every five units increase; *P* = 0.038) remain associated with the PASC phenotype independently from BMI and/or hypertension (**Fig. 2L**). The high levels of fungal translocation during PASC were confirmed using PASC samples from the Rush PASC cohort; these PASC samples were compared to samples from age and sex matched SARS-CoV-2 negative controls (**Fig. 2M**). In the Rush cohort, the majority (74%) of individuals with PASC had β-glucan levels ≥40 pg/ml, whereas only 12% of SARS-CoV-2 negative controls had ≥40 pg/ml of β-glucan (**Fig. 2M**). Together these data suggest that PASC is associated with elevated levels of markers of tight junction permeability (zonulin) and fungal translocation (β-glucan) to the blood.

### Plasma β-glucan levels associate with markers of inflammation during PASC

It is well established that β-glucan can directly induce inflammation following its binding to Dectin-1 on macrophages, monocytes, and dendritic cells. This activates the NF-κB pathway and induces the secretion of pro-inflammatory cytokines (60–62). We, therefore, tested whether β-glucan levels correlated with markers of inflammation, as well as number of symptoms, and overall health/QoL score (**Fig. 3A**). We found a positive correlation between β-glucan levels of inflammatory markers, including TNFα, IL-6, and IP-10 **(Fig. 3A, D-E**). β-glucan levels also associated with and a higher number of symptoms **(Fig. 3A, B)** and a lower overall health/QoL score **(Fig. 3A, C).** The positive correlations between levels of β-glucan and higher IL-6 and TNFα were confirmed in the PASC samples from the Rush Cohort **(Fig. 3F-G**). These data suggest a potential link between fungal translocation and inflammation in individuals with PASC.

**Figure 3.**
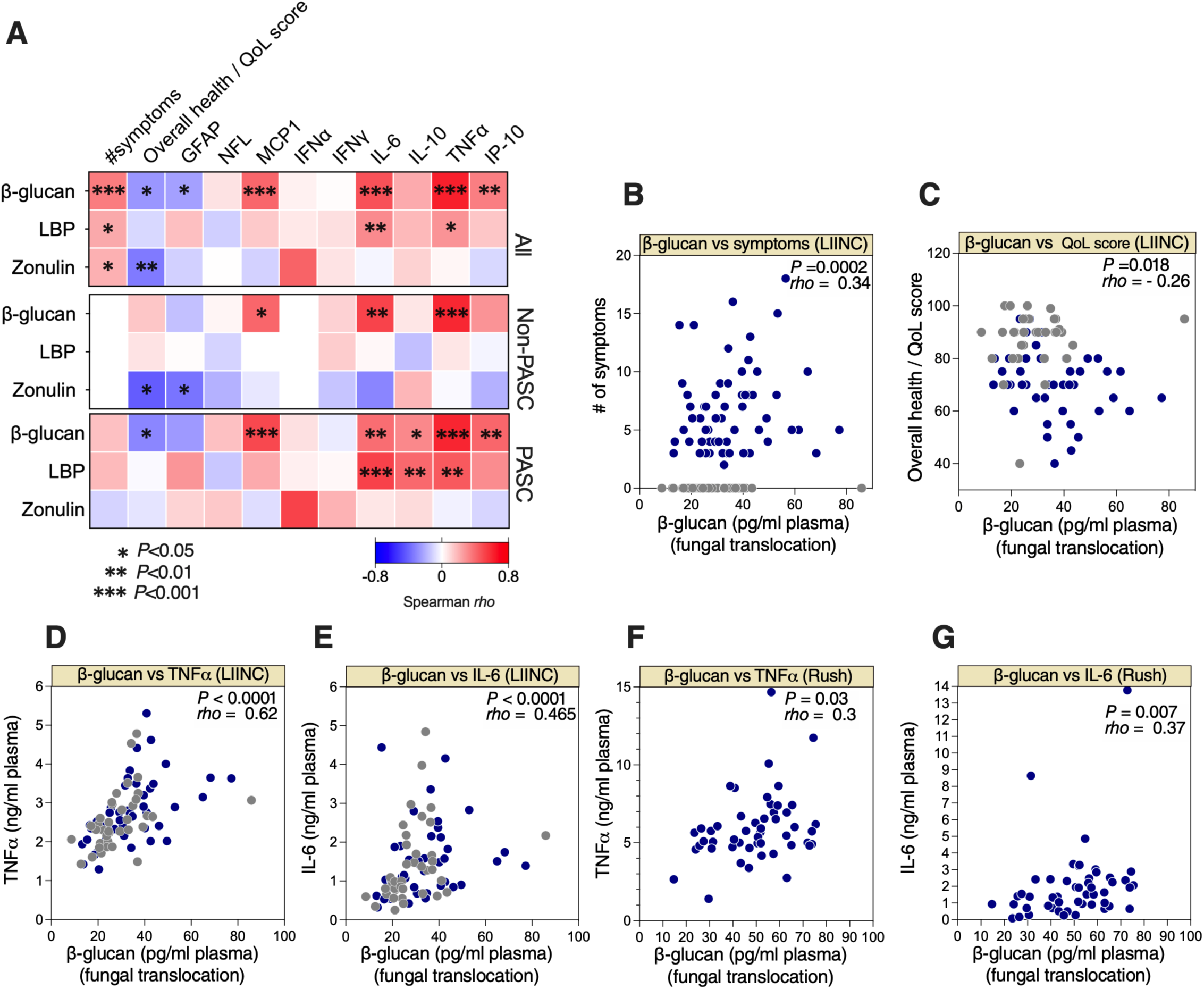
Fungal translocation correlates with inflammation during PASC. **(A)** Three correlation heat-maps showing associations between β-glucan, LBP, or zonulin (in rows) and the number of symptoms during PASC, overall health/quality of life (QoL) score, and plasma levels of several inflammatory markers (in columns) measured in all (n=117; top), non-PASC (n=56; middle), and PASC (n=61; bottom) groups from the UCSF LIINC cohort. The color of the square represents the strength of the Spearman’s rank correlation, with blue shades represent negative correlations and red shades represent positive correlations. * = p< 0.05; ** = p<0.01; *** = p<0.001. **(B-G)** Examples of the correlations between β-glucan and number of symptoms (UCSF LIINC cohort) **(B)**, β-glucan and overall health/QoL score (UCSF LIINC cohort) **(C)**, β-glucan and TNFα or IL-6 (UCSF LIINC cohort) **(D-E)**, β-glucan and TNFα or IL-6 (Rush PASC cohort) **(F-G).** Spearman’s rank correlation tests were used for statistical analysis. Blue = PASC, and grey = non-PASC in B-E.

### Plasma β-glucans from PASC patents activate the NF-κB pathway

The data described thus far suggest that PASC is associated with high plasma levels of β-glucan in a manner linked to higher inflammation. Although modest compared to levels observed during invasive fungal infections, the levels of plasma β-glucan we observed in two cohorts of PASC (Fig. 2-3) may be clinically significant and with a potential to exacerbate a pro-inflammatory state, as suggested by a recent study (57). In that study (not focus on COVID-19), β-glucan levels in plasma ≥40 pg/ml were associated with higher inflammation, fewer ventilator-free days, and worse survival in patients with acute respiratory failure (57). β-glucans are known to induce inflammation by activating the NF-κB pathway following binding to the Dectin-1 receptor (60–62). We, therefore, examined whether there is a mechanistic link between the levels of β-glucans in the plasma of individuals with PASC and the Dectin-1 dependent activation of the NF-κB pathway. For these experiments, we used a dectin-1 receptor reporter cell line to measure β-glucan/Dectin-1 dependent NF-κB signaling. This cell line stably expresses the Dectin-1 receptor and an NF-κB reporter linked to secreted alkaline phosphatase (SEAP) so that dectin-1 receptor stimulation by β-glucan can be measured by quantifying SEAP activity (**Fig. 4A**).

**Figure 4.**
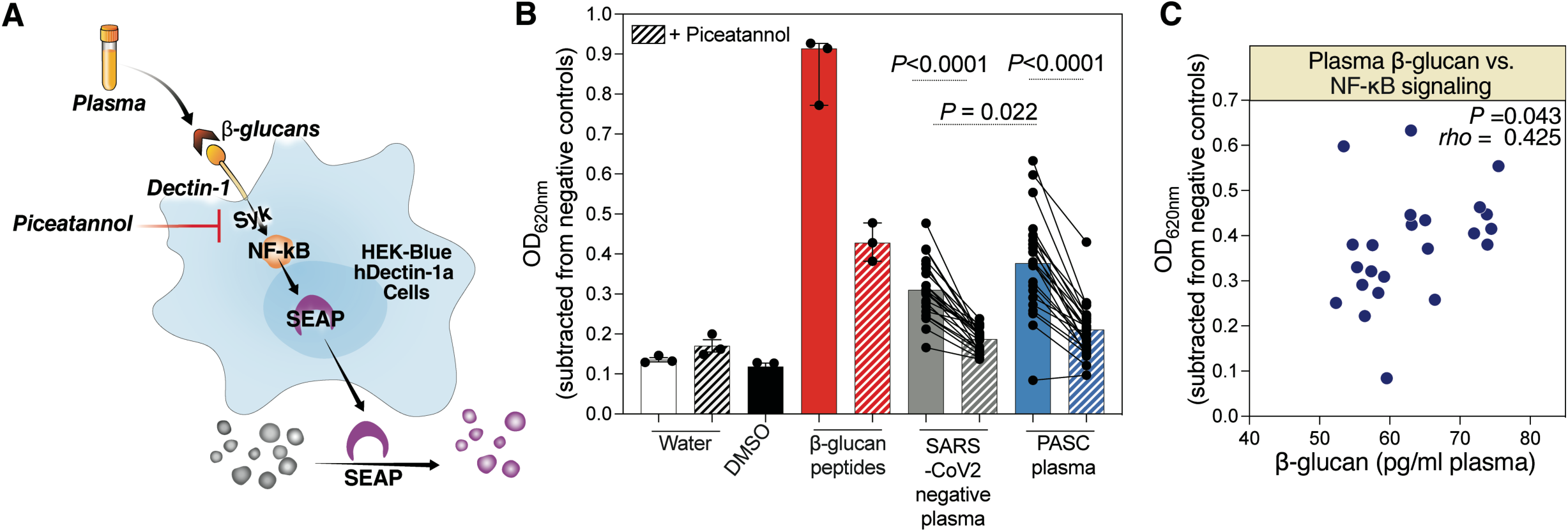
Plasma from individuals with PASC induces NF-kB signaling, which is dampened by the Syk inhibitor Piceatannol. **(A)** A schematic of the dectin-1 receptor reporter cell line. This cell line stably expresses the dectin-1 receptor and an NF-κB reporter linked to secreted alkaline phosphatase (SEAP) so that dectin-1 receptor stimulation by β-glucan can be measured by quantifying SEAP activity. **(B)** The Dectin-1 receptor reporter cells were treated with plasma from individuals with PASC or SARS-CoV-2 negative controls in the presence of absence of the Syk inhibitor, Piceatannol. NF-kB signaling was detected (OD_620nm_) 24 hours later. The comparison between the SARS-CoV-2 negative controls and PASC samples without Piceatannol was performed using Mann–Whitney U test and the comparisons between the conditions with and without Piceatannol were performed using Wilcoxon signed rank tests. **(C)** Spearman’s rank correlation between levels of β-glucan in the plasma during PASC and NF-κB signaling induced by PASC plasma, using the dectin-1 receptor reporter system.

We treated the Dectin-1 receptor reporter cells with 20µl of plasma from volunteers with PASC (a subset of the Rush PASC cohort) or 20µl of plasma from SARS-CoV-2 negative controls. The plasma from the PASC group induced significantly higher levels of NF-kB activation (in a Dectin-1 and β-glucan dependent manner) compared to the plasma from SARS-CoV-2 negative controls (**Fig. 4B**). This β-glucan/Dectin-1 dependent NF-κB signaling was significantly abrogated by the addition of a selective inhibitor to the Syk signaling (Piceatannol) (**Fig. 4B**). Finally, levels of β-glucan in the plasma of those with PASC correlated with a higher ability of these plasma samples to induce β-glucan/Dectin-1 dependent NF-κB signaling using this reporter system (**Fig. 4C**). These data suggest that the levels of β-glucan in the plasma during PASC are capable of inducing immune activation in a manner linked to the activation of the Dectin-1-Syk-NF-κB signaling pathway. Together, these observations suggest a potential mechanism by which fungal translocation may contribute to inflammation during PASC. Importantly, this inflammation can be inhibited using inhibitors to dectin-1/Syk, providing a potential approach to mitigate PASC.

### PASC is associated with elevated levels of host metabolic agonists of the *N*-Methyl-D-aspartate (NMDA) receptors with established neuro-toxic properties

Microbial translocation-mediated inflammation can not only impact biological functions directly, but it also may impact them indirectly by modulating the circulating levels of metabolites derived from interactions between gut microbiota and the host. Many plasma metabolites are biologically active molecules able to regulate cellular processes and immunological functions (63). For example, inflammation-mediated tryptophan catabolism has been associated with the development of several aging- and inflammation-associated diseases during HIV infection (64–68). Severe acute COVID-19 has been associated with a disruption in the levels of several host metabolites such as the metabolites involved in the tryptophan catabolism pathway and S-sulfocysteine (46, 69). We therefore performed an untargeted metabolic analysis (using LC-MS/MS) on the plasma samples from the UCSF LIINC cohort. Within the 117 plasma samples, we identified 169 polar metabolites. We observed a significant (with nominal *P* value <0.05) difference between PASC and non-PASC groups in 12 of these metabolites (six were higher in the PASC compared to non-PASC group and six were lower in the PASC compared to non-PASC group: **Supplementary Table 1**). Untargeted metabolite enrichment analysis of these 12 PASC-associated metabolites showed an enrichment of amino acids and certain amino-acid-related metabolic pathways (**Fig. 5A**).

**Figure 5.**
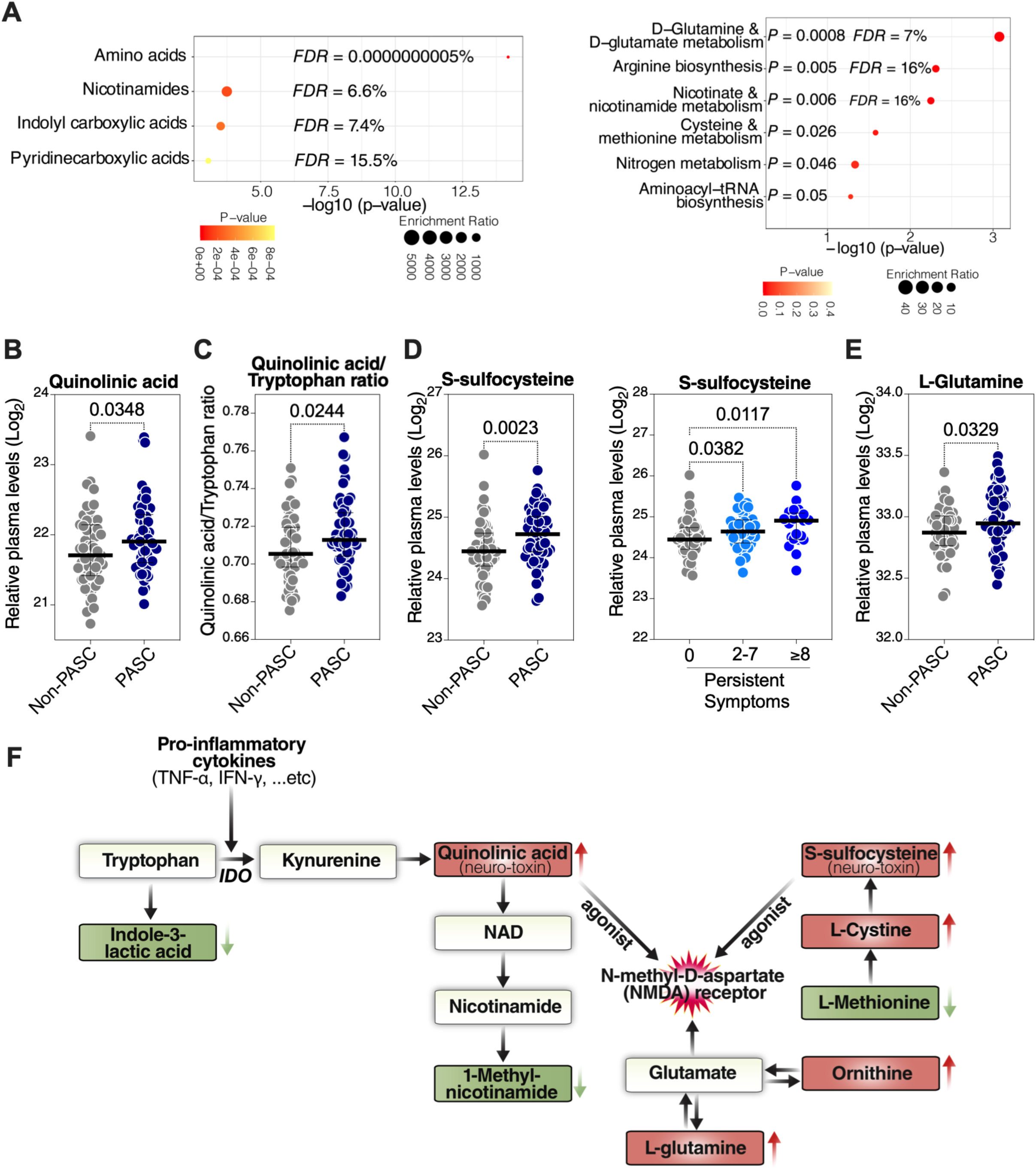
PASC is associated with elevated levels of host metabolic agonists of the *N*-Methyl-D-aspartate receptors with established neuro-toxic properties. **(A)** Unbiased enrichment analyses of the 12 plasma metabolites whose levels differ between PASC and non-PASC groups within the UCSF LIINC cohort. Analysis was performed using MetaboAnalyst 5.0 (http://www.metaboanalyst.ca/). Left image: enrichment of certain classes of metabolites. Right image: enrichment of certain metabolic pathways using the KEGG database. FDR= False Discovery Rate. (**B-E**) Mann–Whitney U comparisons of the plasma levels of quinolinic acid (**B**), quinolinic acid/tryptophan (Q/T) ratio (**C**), S-sulfocysteine (**D**), or L-glutamine **(E**) in PASC and non-PASC groups from the UCSF LIINC cohort. Median and IQR are displayed. (**F**) A model depicting eight plasma metabolites whose levels differ between PASC from non-PASC groups (red is higher in PASC than non-PASC and green is lower in PASC than non-PASC) within the UCSF LIINC cohort and their relationship to both the tryptophan catabolism pathway and the activation of the *N*-Methyl-D-aspartate (NMDA) receptors.

Among the differences between PASC and non-PASC groups were higher levels of quinolinic acid, a downstream product of the tryptophan catabolism pathway, in those with PASC compared to the non-PASC group (**Fig. 5B**). Tryptophan catabolism is commonly indicated by two ratios, the kynurenine to tryptophan (K/T) ratio and the quinolinic acid to tryptophan (Q/T) ratio (70). Although we did not observe a statistically significant difference in the K/T ratio between PASC and non-PASC groups, the Q/T ratio was higher in those with PASC compared to the non-PASC group (**Fig. 5C**). Higher levels of quinolinic acid and a higher Q/T ratio levels have been associated with adverse disease outcomes during chronic HIV infection (64, 70). Quinolinic acid is an established neurotoxin and NMDA receptor agonist (71, 72). Interestingly, other metabolites that also activate NMDA receptors were elevated in the plasma of those with PASC, such as S-sulfocysteine (73) (**Fig. 5D**), and L-glutamine (**Fig. 5E**). Indeed, several of the 12 metabolites that differ between those with and without PASC are involved in pathways related to the activation of NMDA receptors, as shown in the diagram in **Fig. 5F**. Consistent with the neurotoxic ability of quinolinic acid and S-sulfocysteine, higher Q/T and K/T ratios were associated with neuropathy during PASC (**Supplementary Fig. 2**) and higher levels of S-sulfocysteine were associated with neurodegenerative PASC (and the other two PASC phenotypes; **Supplementary Fig. 3**). S-sulfocysteine levels were also associated with particular neurological symptoms during PASC, such as vision problems, fatigue, headache, and dizziness (**Supplementary Fig. 4**). Together, these data indicate that a metabolic signature associated with PASC is compatible with increased tryptophan catabolism and accumulation of metabolites with neuro-toxic properties, conferred by their ability to activate NMDA receptors.

### Plasma metabolomic markers of PASC are associated with higher inflammation and lower overall health

As bioactive molecules, plasma metabolites influence cellular processes and immunological responses. Therefore, we asked whether any of the 12 dysregulated plasma metabolites, as well as Q/T and K/T ratios (as markers of tryptophan catabolism), associated with levels of β-glucan, number of symptoms, health score, or plasma markers of inflammation (**Fig. 6A** shows heat-maps focusing on the correlations between Q/T ratio, K/T ratio, and five elevated metabolites; a complete list of correlations are shown in **Supplementary Table 2**). The most significant associations were between Q/T ratio, quinolinic acid, or K/T ratio, and lower health score (only in the PASC group but not in the non-PASC group; **Fig. 6A-D**). In addition, levels of quinolinic acid (and other elevated metabolites) and Q/T and K/T ratios correlated with higher levels of markers of inflammation (**Fig. 6A**), mainly during PASC. These data further support potential links between disrupted metabolic activities, especially those related tryptophan catabolism and NMDA receptor activation, and both inflammation and disease severity during PASC.

**Figure 6.**
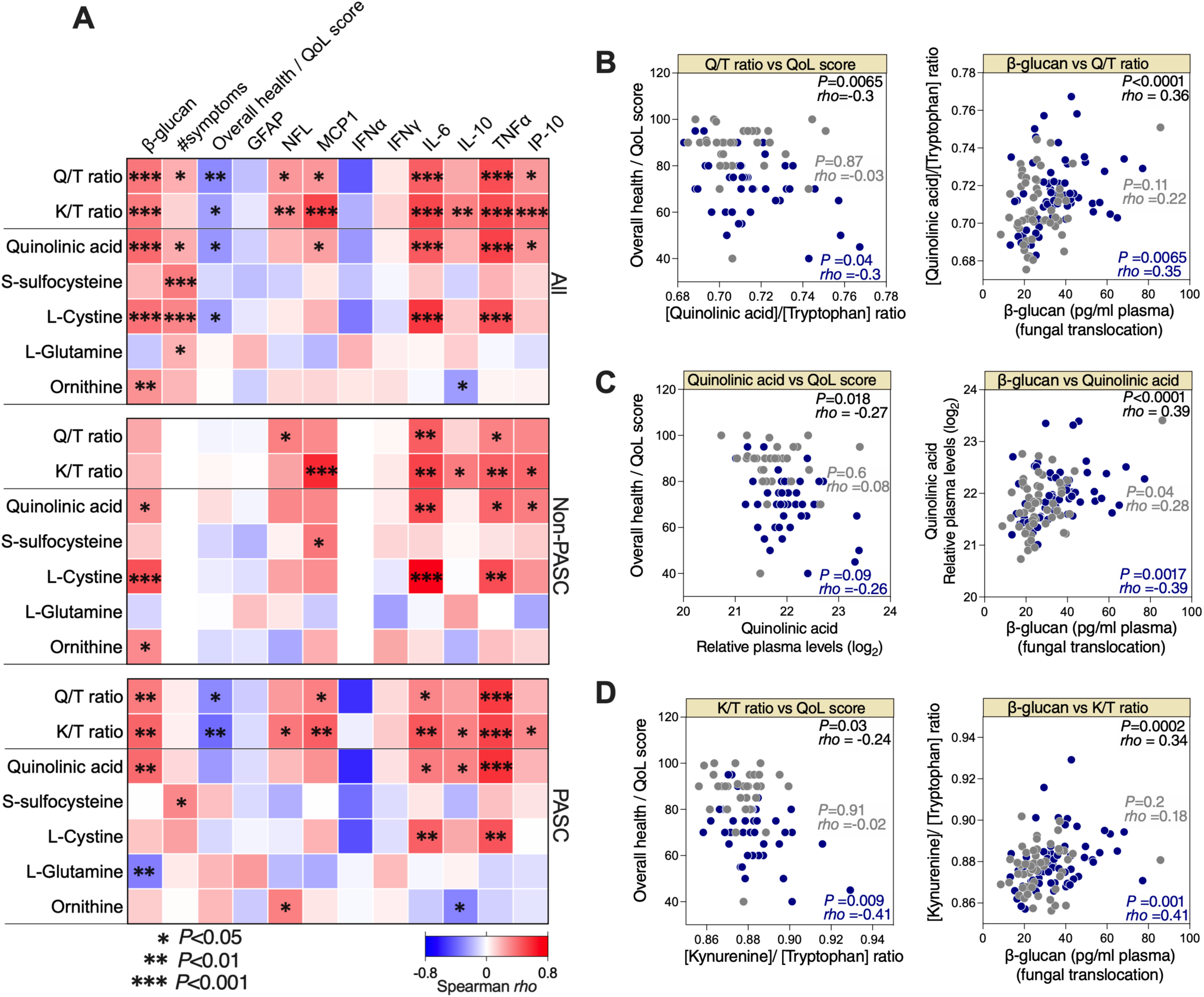
Levels of plasma host metabolites correlate with inflammation during PASC. **(A)** Three correlation heat-maps showing associations between quinolinic acid/tryptophan (Q/T) ratio, kynurenine-to-tryptophan (K/T) ratio, and levels of selected metabolites (in rows) to levels of plasma β-glucan, the number of symptoms during PASC, overall health/quality of life (QoL) score, and plasma levels of several inflammatory markers (in columns) measured in all (n=117; top), non-PASC (n=56; middle), and PASC (n=61; bottom) individuals from the UCSF LIINC cohort. The color of the square represents the strength of the Spearman’s rank correlation, with blue shades representing negative correlations and red shades representing positive correlations. * = p< 0.05; ** = p<0.01; *** = p<0.001. **(B)** Examples of the correlations between Q/T ratio and overall health/QoL score or β-glucan. **(C)** Examples of the correlations between quinolinic acid and overall health/QoL score or β-glucan. **(D)** Examples of the correlations between K/T ratio and overall health/QoL score or β-glucan. Spearman’s rank correlation tests were used for statistical analysis. Blue = PASC, and grey = non-PASC in B-D.

## DISCUSSION

Identifying the potential mechanisms underlying the sustained elevated levels of immune activation and inflammation during PASC is a critical step toward developing tools to prevent or decrease the severity of this syndrome. Our data support a model where a disturbance in tight junction permeability (in the gut and/or lung) can allow fungal translocation to the blood. The fungal β-glucan can then induce the production of pro-inflammatory cytokines by binding to its receptor (Dectin-1) on the surface of myeloid cells and activating the Syk/NF-κB pathway. This activation can be inhibited with the Syk inhibitor, Piceatannol. Our data also suggest that elevation of specific microbiome and inflammation-associated metabolic pathways may contribute to PASC, in particular, pathways with neurotoxic properties due to activation of NMDA receptors (**Fig. 7**). It is unlikely that the elevated levels of fungal translocation and neurotoxic host metabolites are the primary triggers of PASC, as this complex clinical syndrome likely results from the disruption of multiple and probably distinct pathophysiological pathways. However, the robust literature indicating that fungal translocation fuels inflammation and disease severity during long-term complications of other viral infections, such as HIV (28, 31), supports and is consistent with our findings and suggests that fungal translocation may be one of the mechanisms contributing to inflammation during PASC.

**Figure 7.**
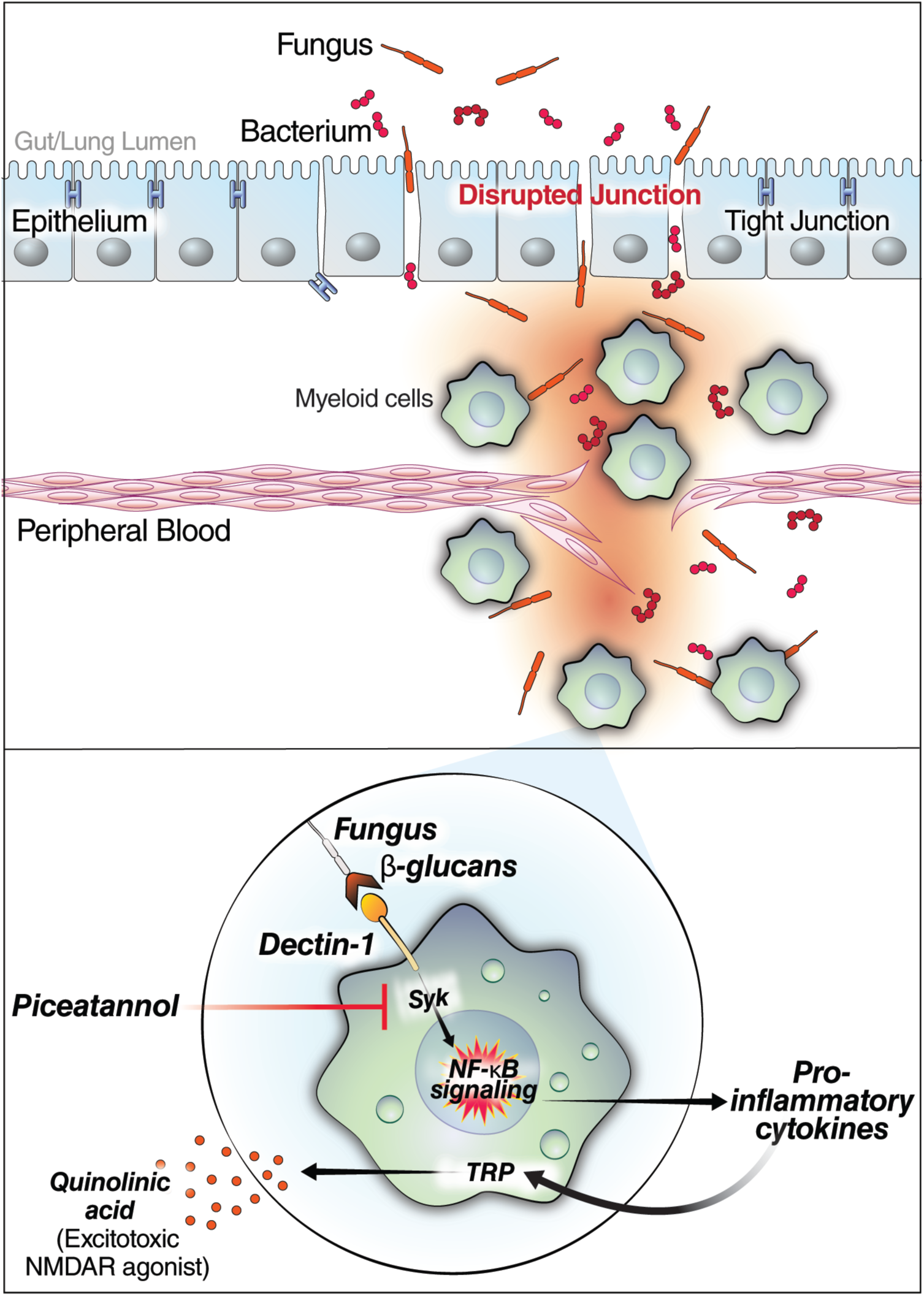
Model of how fungal translocation may contribute to inflammation during PASC. <u>Top</u>: Our data suggest that during PASC there are elevated levels of fungal translocation from the gut and/or lung to the blood (possibly driven by increases in the tight junction permeability driver, zonulin). <u>Bottom</u>: The translocated β-glucans (fungal cell wall polysaccharides) bind to Dectin-1 on myeloid cells to activate the cellular inflammasome via the NF-kB pathway. Blocking the Syk pathway (using Piceatannol) prevents the β-glucan-mediated myeloid inflammation *in vitro* and may prevent it during PASC. Finally, the inflammation (pro-inflammatory cytokines) can activate the tryptophan (TRP) catabolism pathway, and cause other metabolic dysregulations, to induce levels of host metabolic agonists of the *N*-Methyl-D-aspartate receptors (NMDAR; such as quinolinic acid) with established neuro-toxic properties.

Our *in vitro* experiments suggest that the levels of β-glucan in the plasma of individuals experiencing PASC are sufficient to induce NF-κB activation *in vitro* and show that this activation is greater in those with PASC than in SARS-CoV-2 negative controls. As noted above, β-glucan is a biomarker of microbial translocation during chronic viral infections, such as HIV infection, and its levels correlate with inflammation, immune suppression, and the development of HIV-associated co-morbidities (58, 74–78). It can also directly induce inflammation following its binding to Dectin-1 (60–62). Thus, these data suggest a mechanism, Dectin-1-Syk-NF-κB signaling, by which the increased fungal translocation during PASC may contribute to the observed sustained elevated levels of immune activation and inflammation. However, a deeper mechanistic analysis will be needed to identify the degree to which this NF-κB activation contributes to inflammation during PASC. Further analyses could also investigate the possibility that myeloid cells (and other immune cells expressing Dectin-1) from individuals with PASC are less resistant to β-glucan stimulation than cells from individuals without PASC. Together, this deeper analysis could shed light on the causative versus consequential links between fungal translocation, inflammation, and PASC.

Our results support the development of novel strategies to prevent or treat PASC, such as microbial-interaction targeted therapeutics (such as probiotics or metabolites) and/or selective small molecules. For example, small molecules that enhance the epithelial barrier integrity or reduce the detrimental effects of fungal translocation are available, including the zonulin receptor antagonist AT1001 (larazotide acetate); this antagonist decreased the severity and incidence of several inflammation-associated diseases in pre-clinical and clinical studies (79–81) and successfully treated a 17-month-old boy with SARS-CoV-2 associated multisystem inflammatory syndrome in children (MIS-C) who failed anti-inflammatory therapies (82). Also available are the Dectin-1 antagonist, Laminarin, which has been used safely and successfully in mouse models of ischemic stroke (83) and colitis (84), and the Syk signaling inhibitor, Piceatannol, which was used to treat a mouse model of ischemic stroke (83). Our *in vitro* data suggest that Piceatannol may abrogate the β-glucan-mediated inflammation during PASC. These molecules can form a foundation for designing strategies – to be tested pre-clinically as soon as pre-clinical models of PASC are available – to prevent PASC and its long-term complications in individuals recovering from SARS-CoV-2 and/or other similar post-acute infection syndromes.

Aging, and several aging-associating diseases, and even other chronic viral infections, have been associated with a breakdown of homeostasis between the gut and its microbiome (85–87). For example, aging itself changes the composition of the gut microbiota (88–92), leading to microbial translocation, which triggers chronic inflammation (93–95). Aging-associated diseases such as cancer, diabetes, and Alzheimer’s are associated with specific gut microbial signatures (96–106). Chronic HIV infection is associated with a state of gut microbial translocation, which is thought to be a major cause of inflammation (107–115). Even with antiretroviral therapy, the damage to the epithelial barrier caused by HIV is never fully repaired, allowing microbial translocation and inflammation to continue (116–118). Our data raise the question of whether the pre-existing state of microbial translocation and chronic inflammation during these conditions might make individuals living with them more prone to PASC. In this analysis, we included a only small number of HIV-infected individuals (we only analyzed a subset of UCSF LIINC cohort); however, a recent study focused on LIINC participants with HIV suggested that HIV infection is indeed a risk factor for developing PASC (119). Whether the pre-existing microbial translocation and chronic inflammation during HIV infection (107–118), post surviving Ebola disease (120), and/or during other aging-related conditions (96–106) contribute to PASC warrants further investigation.

Our metabolic analysis suggests that PASC is associated with elevated levels of several metabolites with known neurotoxic properties that are linked to activation of the NMDA receptor, such as quinolinic acid and S-sulfocysteine. The overactivation of NMDA receptors can lead to excitotoxicity and has been associated with the development of several neurodegenerative disorders including epilepsy, Parkinson’s, Alzheimer’s, and Huntington’s diseases (121–125). Whether an overactivation of NMDA receptors due to dysregulation of host metabolites contributes to neuropathology during PASC warrants investigation. A demonstration that NMDA receptor agonists contribute to PASC symptoms could have several clinical implications. For example, NMDA receptor antagonists (such as memantine) have been used to block excessive, excitotoxic activity resulting from the overactivation of the NMDA receptors during Alzheimer’s disease (122–126), and have been evaluated for treating other neurodegenerative disorders (127–129). Whether memantine or other NMDA receptor antagonists can be used to prevent or treat PASC-associated neurologic manifestations could then be explored.

This study has several limitations. The source of β-glucan in the plasma is not clear. Fungal translocation during illness can occur from both the gut and the lung (57). Examining the contribution of the gut microbiome (using stool samples and intestinal biopsies) and lung microbiome (using sputum and/or bronchoalveolar lavage) will be needed to determine the source of the fungal translocation during PASC and the fungal species contributing to it. It will also be important to examine fungal translocation and host metabolites in longitudinal samples (from different body fluids including cerebrospinal fluid) to determine whether, and for how long, elevated levels of microbial translocation persist after acute COVID-19. Another possibility to the persistent dysregulation is that the intestinal barrier’s resilience to common intestinal disruptors (such as excessive alcohol or oxidative stress) is lower during PASC, making these individuals more vulnerable to frequent common disruptors, which then can lead to the translocation of microbes that cause systemic inflammation. Finally, correcting for additional potential confounders will require validating our results in larger independent cohorts from varying geographic and demographic settings. Despite these caveats, this study, which is exploratory in nature, sheds light on the potential role of microbial translocation and dysregulation of host metabolic pathways related to NMDA receptor activation in the pathophysiology of PASC. By understanding these potential underpinnings of PASC, this work may serve to identify biomarkers for PASC risk stratification and build a foundation for developing strategies to prevent or reduce the severity of inflammation during PASC.

## METHODS

### Study cohorts

Primary analyses were performed using cross-sectional plasma samples from 117 individuals with prior nucleic acid-confirmed SARS-CoV-2 infection (a subset of the UCSF LIINC cohort, described in detail elsewhere (19)) collected 90-160 days after the first positive SARS-CoV-2 qPCR result; prior work has not identified persistent virus in saliva of these individuals at the time of sampling (130). These participants were divided into two groups based upon responses to a standardized symptom assessment at the time of plasma collection: 56 individuals with no COVID-19 attributed symptoms (non-PASC) and 61 with ≥2 COVID-19-attributed symptoms (PASC; **Table 1**); individuals reporting a single symptom were not included. Validation analyses and *in vitro* experiments were performed using cross-sectional plasma samples from 50 individuals with COVID-19 experiencing PASC symptoms 3-4 months after acute COVID-19 (a subset of the Rush PASC cohort) and cross-sectional plasma samples from 50 SARS-CoV-2 negative controls (matched for age and gender to the Rush PASC samples; **Table 1**).

### Ethics

All research protocols were approved by the institutional review board (IRB) at University of California San Francisco (UCSF), Rush University, and The Wistar Institute. All human experimentation was conducted in accordance with the guidelines of the US Department of Health and Human Services and those of the authors’ institutions.

### Symptoms and quality of life score evaluation

Participants in the LIINC study underwent clinical assessment at the time of biospecimen collection. Volunteers completed an interviewer-administered questionnaire querying the presence of 32 possible COVID-19-attributed symptoms, quality of life, and overall health status. The questionnaire was derived from several validated instruments (131, 132) as well as the US Centers for Disease Control list of COVID-19 symptoms. Importantly, a symptom must be described as new or worsened since the diagnosis of SARS-CoV-2 infection to be recorded as “present”; symptoms which existed prior to SARS-CoV-2 infection or were unchanged following infection are not counted. The utility of this instrument in measuring participants longitudinally has been described (19).

### Measurement of plasma markers of tight junction permeability and microbial translocation

Plasma levels of soluble CD14 (sCD14), soluble CD163 (sCD163), and LPS Binding Protein (LBP) were quantified using DuoSet ELISA kits (R&D Systems; catalog # DY383-05, # DY1607-05, and # DY870-05, respectively). The plasma level of Zonulin was measured using an ELISA kit from MyBiosorce (catalog # MBS706368). β-D-glucan detection in plasma was performed using Limulus Amebocyte Lysate (LAL) assay (Glucatell Kit, CapeCod; catalog # GT003).

### Measurement of plasma markers of inflammation

A targeted panel of markers of inflammation in plasma were measured using the HD-X Simoa platform. These markers were selected based upon their importance in acute SARS-CoV-2 infection and included: Cytokine 3-PlexA (IL-6, IL-10, TNFα), interferon-gamma, IP-10, and MCP-1. Levels of these markers in a subset of LIINC participants have been previously reported (22, 24). The IL6 and TNFα measurements of the PASC samples from the Rush PASC cohorts were performed using customized MSD U PLEX multiplex assay (Meso Scale Diagnostic catalog# K15067L-2).

### Untargeted measurement of Plasma metabolites

Metabolomics analysis was performed as described previously (133, 134). Briefly, polar metabolites were extracted with 80% methanol. A quality control (QC) sample was generated by pooling equal volumes of all samples and was injected periodically during the sample sequence. Liquid chromatography-mass spectrometry (LC-MS) was performed on a Thermo Scientific Q Exactive HF-X mass spectrometer with HESI II probe and Vanquish Horizon UHPLC system. Hydrophilic interaction liquid chromatography (HILIC) was performed at 0.2 ml/min on a ZIC-pHILIC column (2.1 mm × 150 mm, EMD Millipore) at 45°C. All samples were analyzed by full MS with polarity switching, and the QC sample was also analyzed by data-dependent MS/MS with separate runs for positive and negative polarities. Raw data were analyzed using Compound Discoverer 3.1 (ThermoFisher Scientific). Metabolites were identified either by accurate mass and retention time using an in-house database generated from pure standards or by querying the mzCloud database (www.mzCloud.org) with MS/MS spectral data and selecting matches with 50 or greater scores. Metabolite quantification used integrated peak areas from full MS runs. These values were corrected based on the periodic QC runs and normalized to the total signal from identified metabolites in each sample.

### Reporter assay for Dectin-1 activation by β-glucan

HEK-Blue hDectin-1a cells (Invivogen; catalog #hkb-hdect1a) was maintained in growth medium containing DMEM (4.5 g/l glucose), 10% fetal bovine serum (FBS), 100 U/ml penicillin, 100 µg/ml streptomycin, 100 µg/ml Normocin, 2 mM L-glutamine, 1 µg/ml puromycin, and 1X HEK Blue CLR Selection (Invivogen; catalog #hb-csm). This cell line expresses the Dectin-1a isoform and genes involved in the Dectin-1/NF-κB/ Embryonic Alkaline Phosphatase (SEAP) signaling pathway. On the assay day, 180 µl/well of cells at a concentration of 2.8 × 10^5^ cells/ml in HEK-Blue detection media (Invivogen; catalog #hb-det2) were plated in 96 well tissue-culture plate. Plasma samples (20 µl) were added to each well with and without Piceatannol (250nM). The plates were then the incubated at 37°C and 5% CO2 for 24 hrs. Levels of SEAP were monitored and measured spectrophotometrically at 620nm. As controls, 20µl of water, Piceatannol (250nM), 2µl of DMSO (Piceatannol solvent), 20 µl of 10 µg/ml β-glucan peptides (Invivogen; catalog #tlrl-bgp), 20 µl of β-glucan peptides + Piceatannol (250nM) were added in separated wells.

### Statistical analysis

Mann–Whitney U tests were used in the analyses of Figures 1A-C, 2A-F, 2H, 2J, 4B (in the comparisons between the SARS-CoV-2 negative controls and PASC samples without Piceatannol), and 5BE. Fisher’s exact tests were used in the analysis in Figure 1D. Spearman’s rank correlations were used in the analyses in Figures 2G, 3, and 6. A multivariate logistic regression model adjusting for BMI and hypertension was used for each marker in the analysis in Figure 2I. Wilcoxon signed rank tests were used in the analysis in Figure 4B (in the comparisons between the conditions with and without Piceatannol). Kruskal-Wallis tests were used in the analyses in Supplementary Figures 1, 2, and 3. All statistical analyses were performed in R and Prism 9.0 (GraphPad).

## AUTHOR CONTRIBUTIONS

M.A-M conceived and designed the study. L.B.G, J.K, K.Y.H, A.R.G, and H-Y.T carried out the experiments. B.C.Y, A.C, J.W.W, and C.J.P measured the inflammatory markers. M.J.P, G.K, N.F.Z, H.R, G.M, F.B, R.A.B, J.N.M, R.H, S.L, J.D.K, H.W, J.N.M, A.K, A.L, S.G.D, and T.J.H screened and selected study participants, supervised all clinical aspects of the study, and collected and interpreted the clinical data. J.D and Q.L analyzed the data. L.B.G. and M.A-M wrote the manuscript, and all authors edited it.

## ACKNOWLEDGMENTS

This study is supported by a Campbell Foundation grant to M.A-M, Commonwealth of Pennsylvania – COVID-19 funding to M.A-M, NIH R01AA029859 to A.K and M.A-M, a supplement to the NIH R01 DK123733 to M.A-M, A.L, and A.K, a supplement to the NIH R24 AA026801 to A.K. The Wistar Proteomics and Metabolomics Shared Resource is supported in part by NIH Cancer Center Support Grant CA010815. The Thermo Q-Exactive HF-X mass spectrometer was purchased with NIH grant S10 OD023586. The LIINC study is supported by NIH/NIAID 3R01AI141003-03S1 and NIH/NIAID R01AI158013. The Rush University Medical Center Post-COVID-19 Clinic and Biorepository is supported by an Abbott Laboratories grant to H.W. M.J.P is supported on K23AI157875. J.D.K is supported on K23AI135037. We acknowledge the contributions of the UCSF Clinical and Translational Science Unit, Core Immunology Laboratory, and AIDS Specimen Bank. We would like to thank Rachel E. Locke, Ph.D., for providing comments.

## SUPPLEMENTARY MATERIAL

**Supplementary Figure 1.**
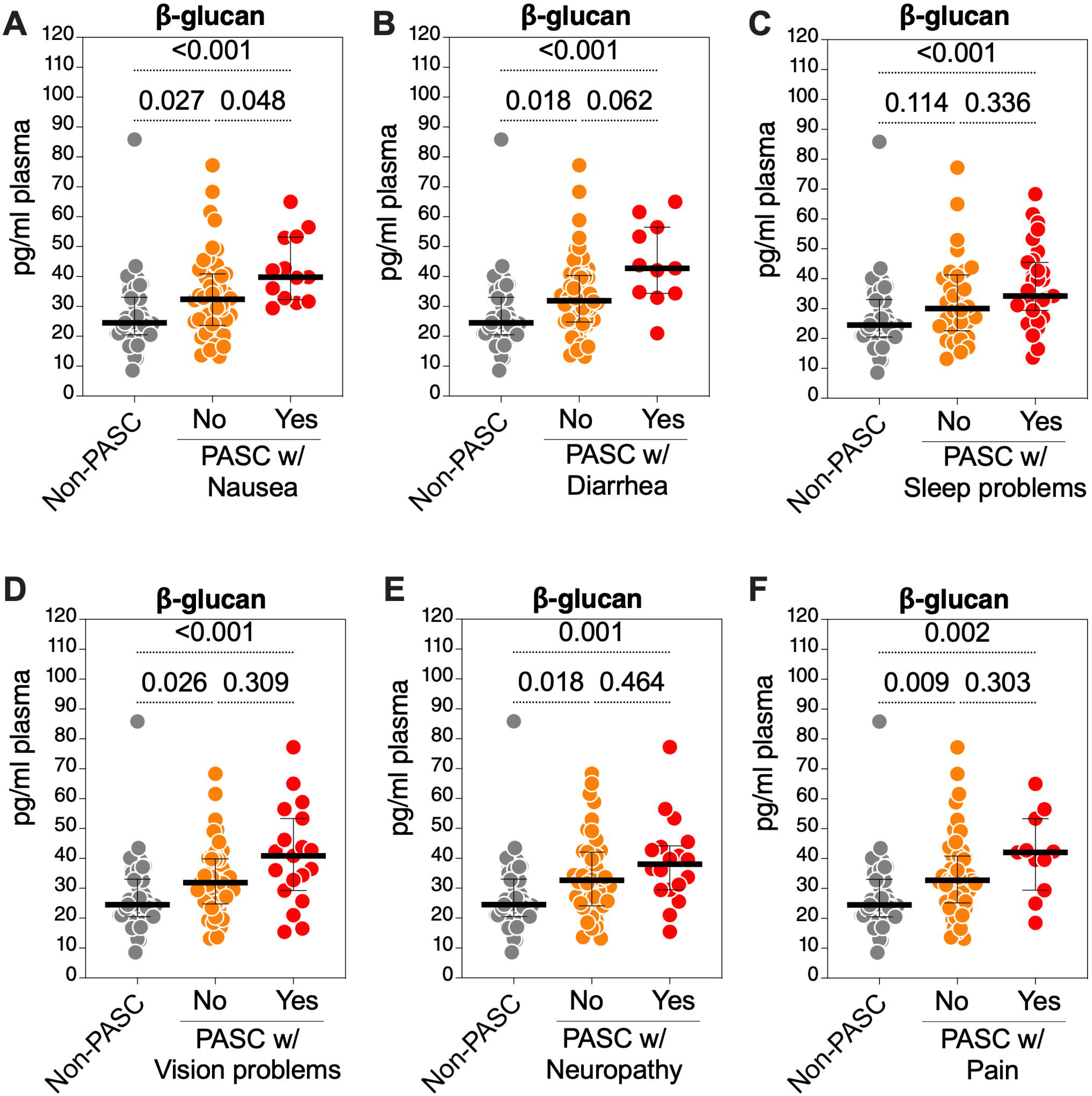
β-glucan levels in plasma correlate with certain PASC symptoms. Kruskal-Wallis tests comparing β-glucan levels in non-PASC and PASC groups (from the UCSF LIINC cohort) with or without **(A)** nausea, **(B)** diarrhea, **(C)** vision problems, **(D)** sleep problems, **(E)** neuropathy, or **(F)** pain. Median and IQR are displayed.

**Supplementary Figure 2.**
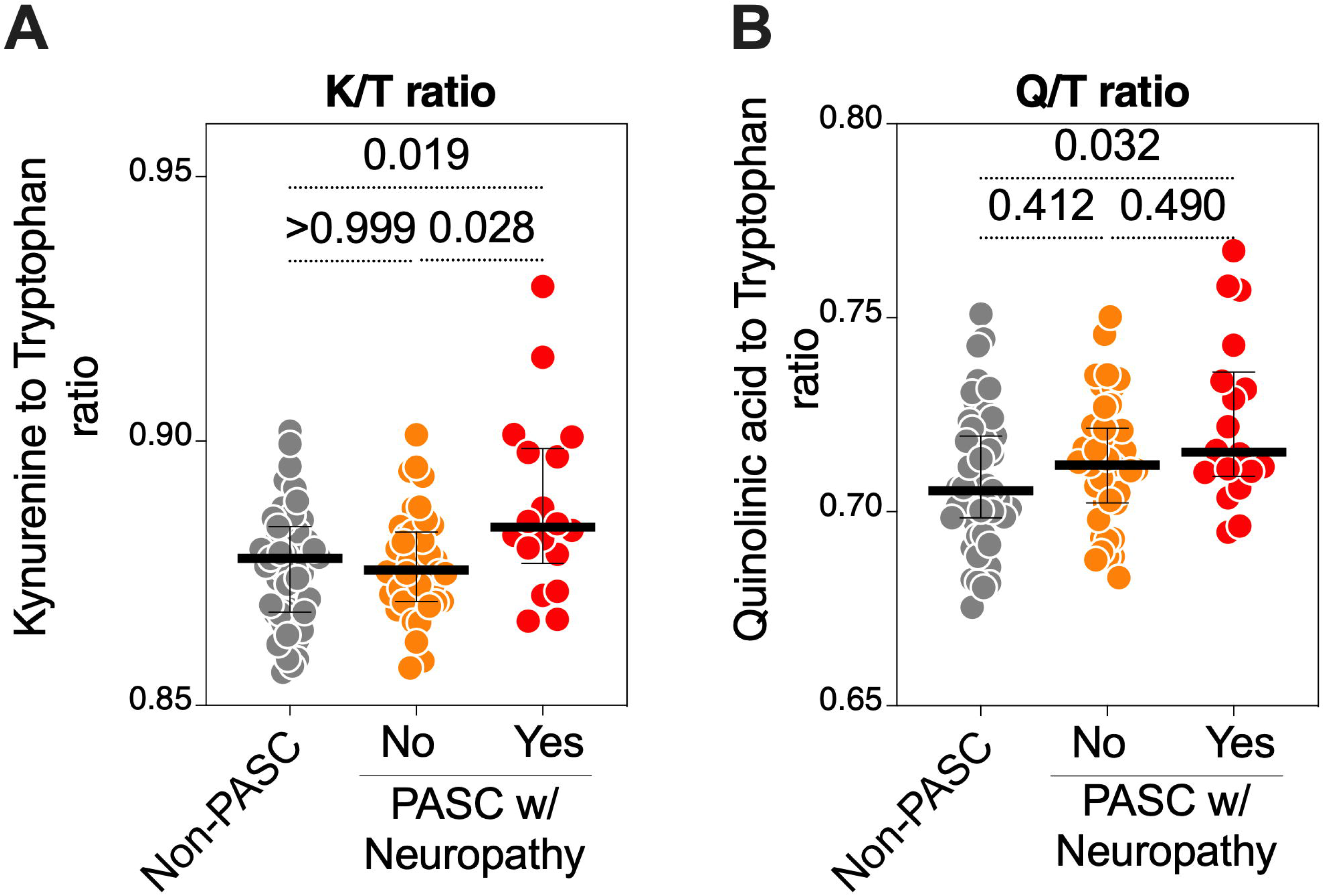
K/T and Q/T ratios correlate with neuropathy during PASC. Kruskal-Wallis tests comparing **(A)** K/T ratio and **(B)** Q/T ratio between non-PASC and PASC groups (from the UCSF LIINC cohort) with or without neuropathy symptoms. Median and IQR are displayed.

**Supplementary Figure 3.**
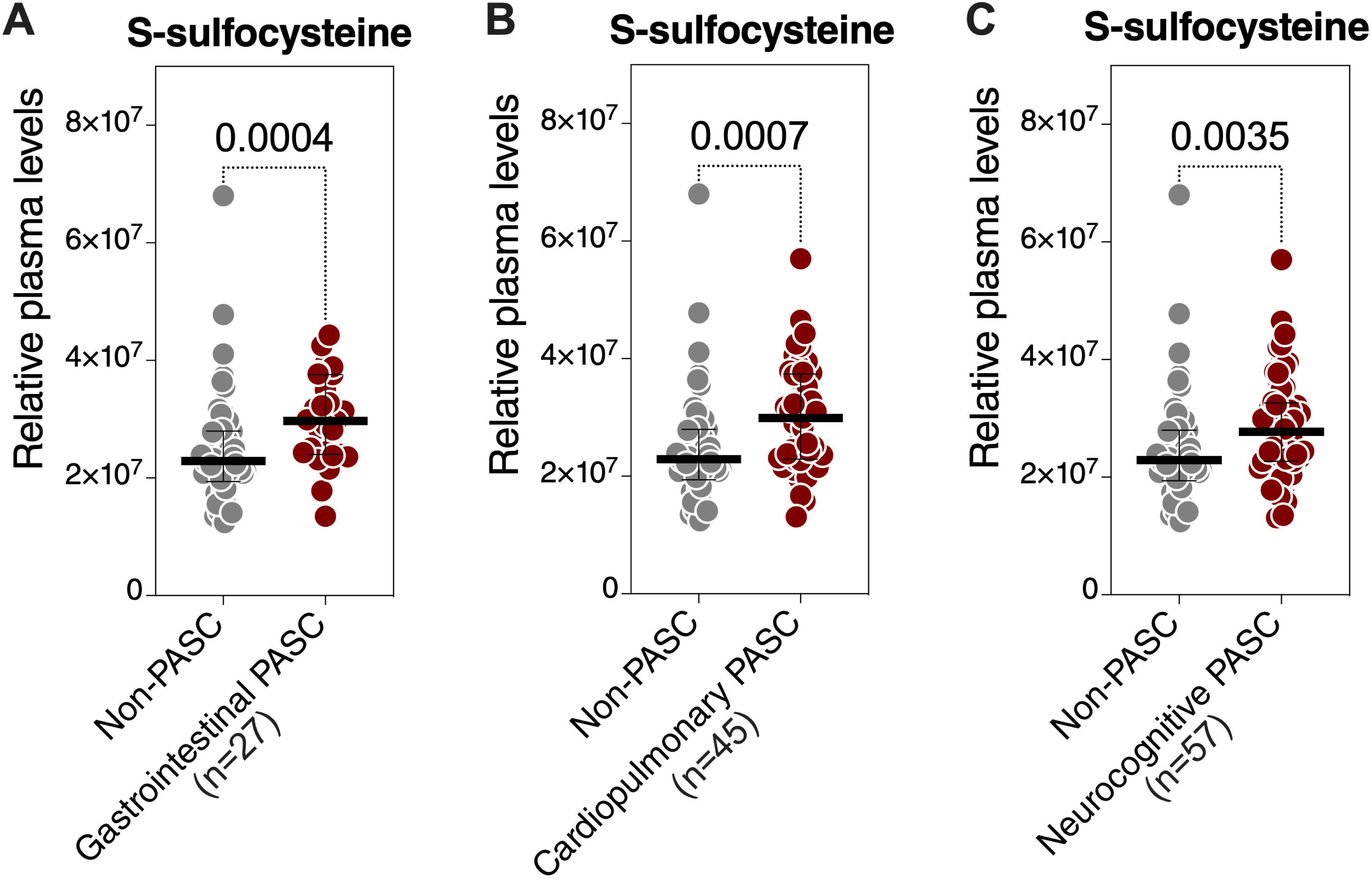
Levels of plasma S-sulfocysteine are higher in individuals experiencing three PASC phenotypes, based on clinical symptom clusters, compared to those without PASC. PASC was divided to three phenotypes based on clinical symptom clusters defined as having at least one symptom in the cluster – gastrointestinal (nausea, diarrhea, loss of appetite, abdominal pain, vomiting), cardiopulmonary (cough, dyspnea, chest pain, palpitations), and neurocognitive (headache, concentration problems, dizziness, balance problems, neuropathy, vision problems). S-sulfocysteine levels were higher in individuals experiencing each of the three PASC symptom clusters compared to individuals who are not experiencing PASC. Mann–Whitney U tests. Median and IQR are displayed.

**Supplementary Figure 4.**
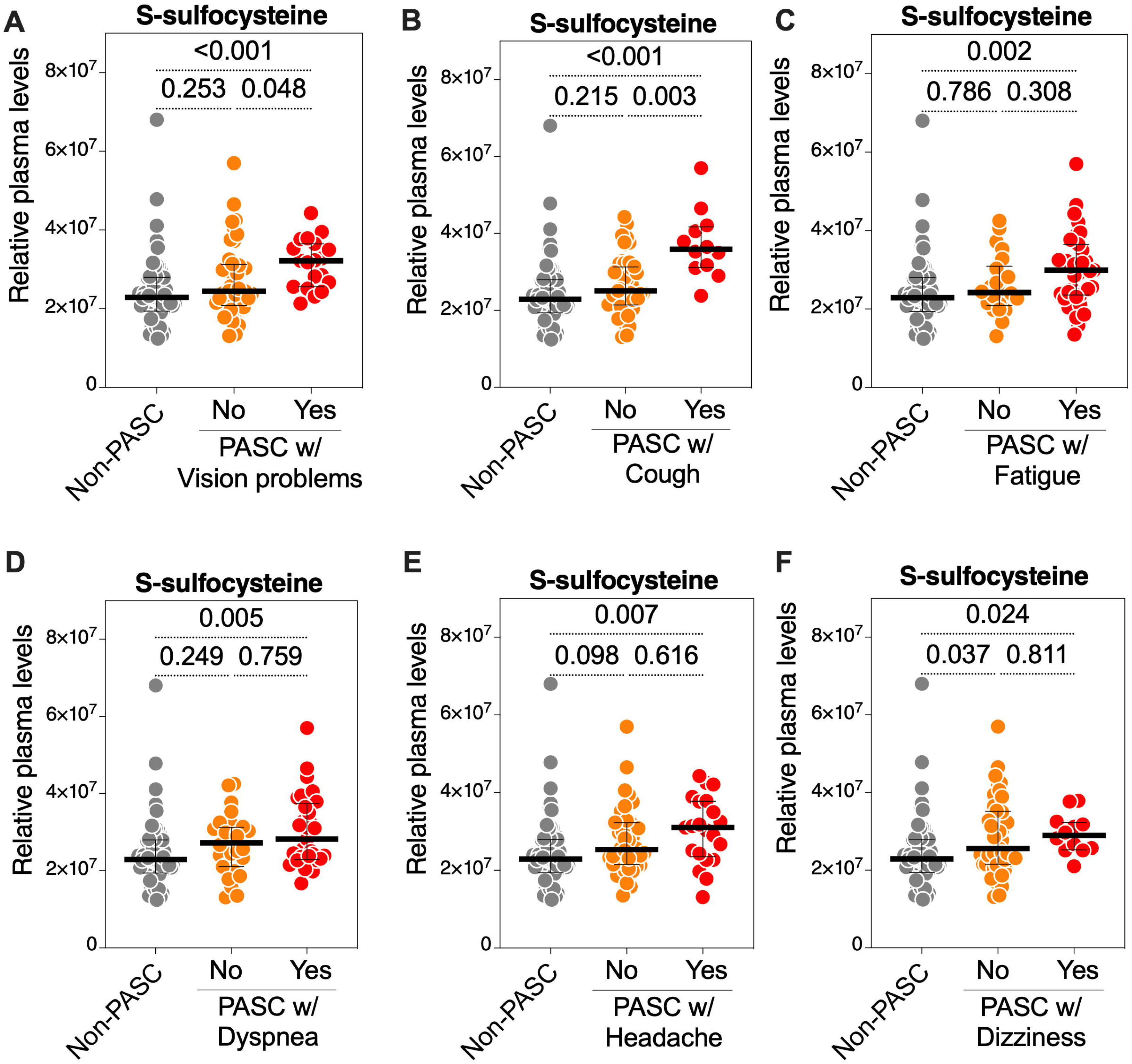
S-sulfocysteine levels in plasma correlate with certain PASC symptoms. Kruskal-Wallis tests comparing S-sulfocysteine levels in non-PASC and PASC groups (from the UCSF LIINC cohort) with or without **(A)** vision problems, **(B)** fatigue, **(C)** headache, **(D)** dizziness, **(E)** cough, or **(F)** dyspnea. Median and IQR are displayed.

**Supplementary Table 1.**
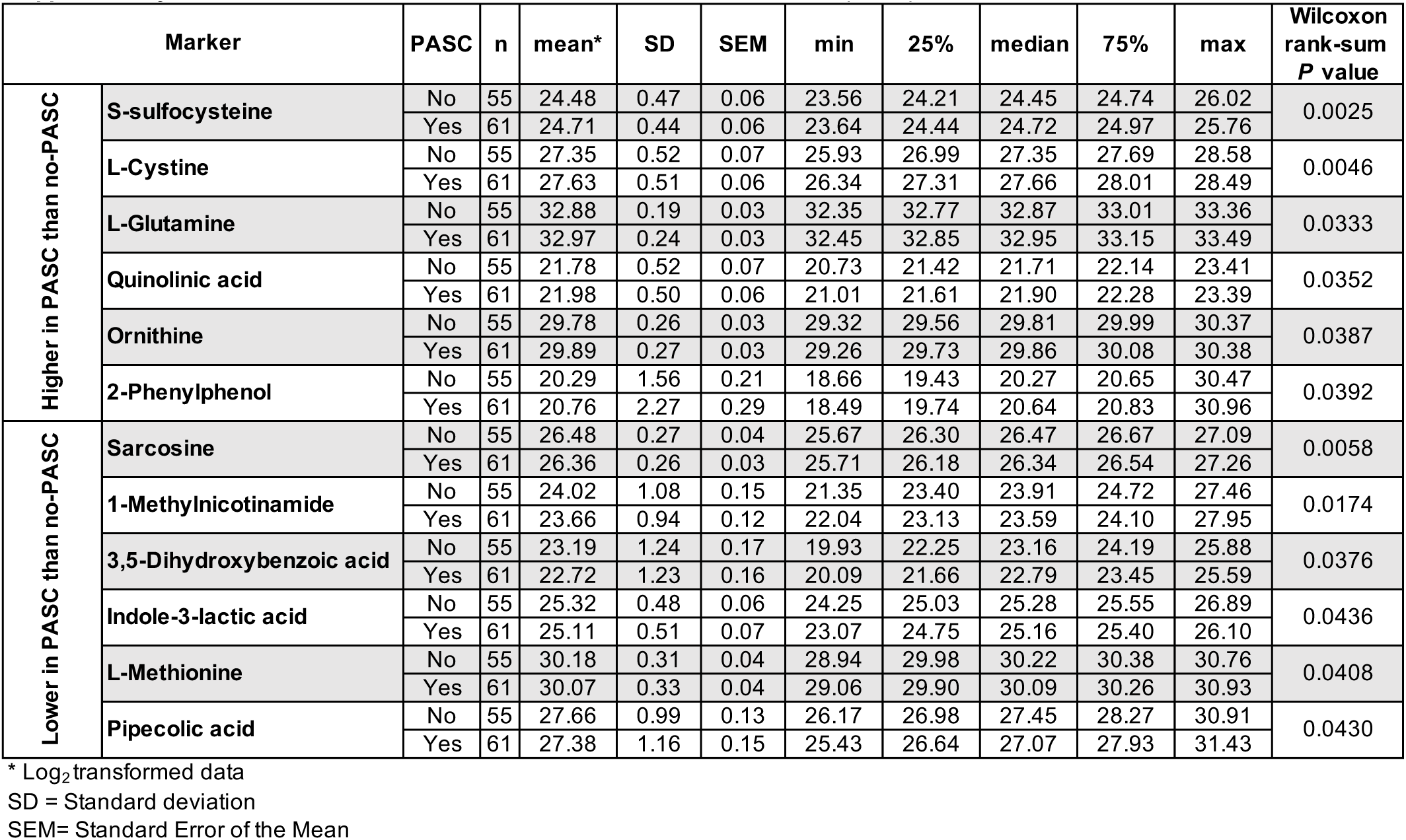
Plasma metabolites differ between PASC and non-PASC participants in the LIINC cohort.

**Supplementary Table 2.**
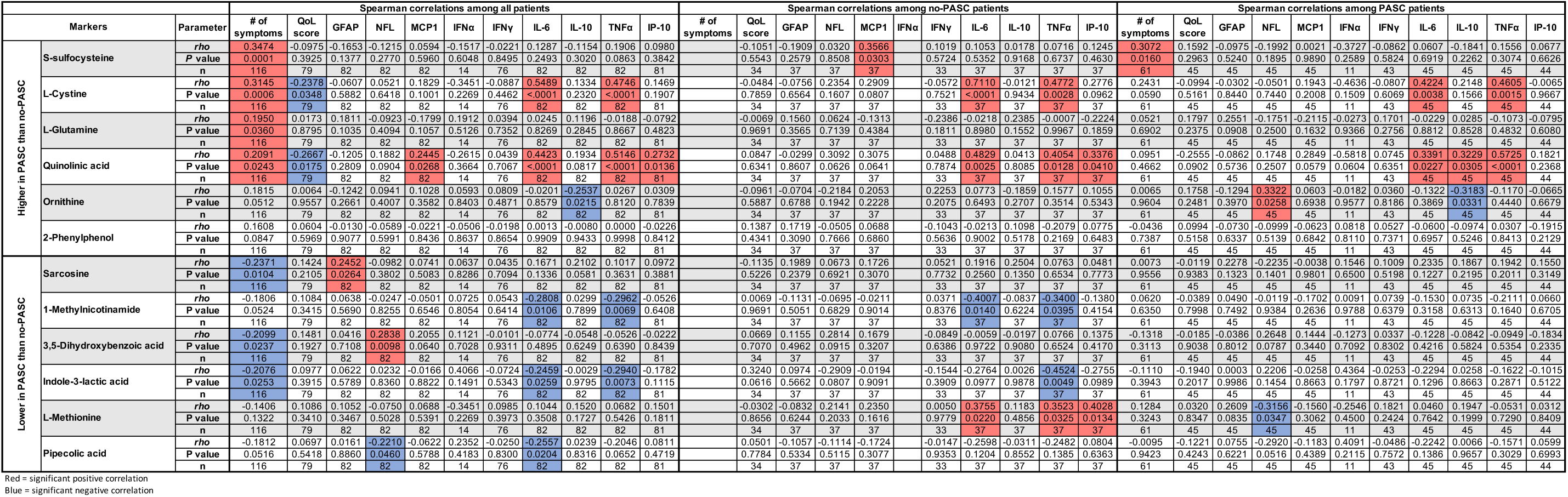
Correlations between metabolites and clinical and inflammatory markers in the LIINC cohort.

